# Mechanistic target of rapamycin complex 1 (mTORC1) activity occurs predominantly in the periphery of human skeletal muscle fibers, in close proximity to focal adhesion complexes, following anabolic stimuli

**DOI:** 10.1101/2021.06.22.449494

**Authors:** Nathan Hodson, Michael Mazzulla, Dinesh Kumbhare, Daniel R. Moore

## Abstract

Following anabolic stimuli (e.g. mechanical loading and/or amino acid provision) the mechanistic target of rapamycin complex 1 (mTORC1), a master regulator of protein synthesis, translocates toward the cell periphery. However, it is unknown if mTORC1 activity occurs prior to or following this translocation. We therefore aimed to determine the cellular location of mTORC1 activity in human skeletal muscle following anabolic stimuli. Fourteen young, healthy males either ingested a protein-carbohydrate beverage (0.25g/kg protein, 0.75g/kg carbohydrate) alone (n=7, 23±5yrs, 76.8±3.6kg, 13.6±3.8%BF, FED) or following a whole-body resistance exercise bout (n=7, 22±2yrs, 78.1±3.6kg, 12.2±4.9%BF, EXFED). *Vastus lateralis* muscle biopsies were obtained at rest (PRE) and 120 and 300min following anabolic stimuli. The spatial regulation of mTORC1 activity was assessed through immunofluorescent staining of p-RPS6^Ser240/244^, an mTORC1-specific phosphorylation event. p-RPS6^Ser240/244^ measured by immunofluorescent staining or immunoblot was positively correlated (r=0.76, p<0.001). Peripheral staining intensity of p-RPS6^Ser240/244^ increased above PRE in both FED and EXFED at 120min (~54% and ~138% respectively, p<0.05) but was greater in EXFED at both post-stimuli time points (p<0.05). The peripheral-central ratio of p-RPS6^240/244^ staining was displayed a similar pattern, suggesting mTORC1 activity occurs predominantly in the periphery of fibers. Moreover, p-RPS6^Ser240/244^ intensity within paxillin-positive regions, a marker of focal adhesion complexes, was elevated at 120min irrespective of stimulus (p=0.006) before returning to PRE at 300min. These data confirm that mTORC1 activity occurs in the region of human muscle fibers to which mTORC1 translocates following anabolic stimuli and identifies focal adhesion complexes as a potential site of mTORC1 activation in vivo.

## Introduction

Skeletal muscle size is governed by net protein balance (NPB), the algebraic difference between muscle protein synthesis (MPS) and breakdown (MPB). Of these two components, MPS is the most sensitive, drastically elevating in response to anabolic stimuli such as amino acid ingestion or mechanical loading (1, 2). At the molecular level, MPS is believed to be primarily regulated by mechanistic target of rapamycin complex 1 (mTORC1), an evolutionarily conserved serine/threonine kinase, whose downstream targets are implicated in the control of translation initiation and elongation and ribosomal RNA synthesis (3). Indeed, the ingestion of rapamycin, an mTORC1 inhibitor, completely abolishes post-exercise/feeding induced elevations in MPS in humans (4, 5). Moreover, mTORC1 also inhibits MPB via the inhibition of autophagophore biogenesis and nucleation (6, 7). mTORC1 is therefore an integral regulator of NPB in skeletal muscle and as such research has focused on elucidating the mechanisms of mTORC1 activation in order to identify interventions to promote MPS and subsequent muscle growth in compromised populations (e.g. older or clinical populations).

Initially, the principal source of data regarding mTORC1 activation was gleaned from *in vitro* investigations utilizing HeLa and/or HEK293 cell lines (8–11). Here, the lysosome was identified as the site of mTORC1 activation due to its abundance of amino acids within its lumen, and the presence of direct mTORC1 activators, Ras homolog enriched in brain (Rheb) and phosphatidic acid, on the lysosomal membrane (12, 13). In response to amino acid provision, mTORC1 is recruited to the lysosomal membrane, from the cytosol, through an ‘inside-out’ mechanism involving the vATPase-Ragulator-Rag protein axis to become active (8–10). In contrast, following growth factor administration, mTORC1 activity is enhanced through removal of tuberous sclerosis complex proteins (TSC1/2) from their association with Rheb allowing Rheb to convert to its active guanosine tri-phosphate loaded state (14). In rodent skeletal muscle, mechanical stimulation was shown to activate mTORC1 by a similar mechanism, reducing TSC1/2-Rheb colocalization and enhancing mTORC1 enrichment at the lysosomal membrane (15). Although lysosomal targeting of this kinase was originally believed to be the mechanism of mTORC1 activation in human skeletal muscle, recent investigations have not specifically supported this. For example, in human skeletal muscle amino acid ingestion, mechanical loading or a combination has little effect on mTORC1-lysosomal colocalization, which may be related to upregulation of autophagy and other catabolic systems that maintain intralysosomal amino acid concentrations preserve mTORC1 localization at the lysosomal surface (16–18). Instead, anabolic stimuli initiate the translocation of mTORC1-lysosome complexes toward the cell periphery where mTORC1 becomes in close proximity to upstream activators (e.g. Akt), downstream substrates (e.g. translation initiation factors) and the vasculature (entry site of extracellular amino acids) (16–21). These findings have also been reinforced by *in vitro* evidence where the disruption of lysosomal movement impairs mTORC1 activation in response to amino acids or growth factors (18). However, although mTORC1-lysosomal translocation and mTORC1 kinase activity appear related in human skeletal muscle, it is currently unknown whether mTORC1 activity occurs prior to (i.e. in central, cytosolic regions) or following this translocation (i.e. in peripheral regions).

A recent *in vitro* investigation has also identified focal adhesion complexes as an important site in mTORC1 regulation. These complexes are enriched with growth factor receptors, amino acid transporters and integrins (22) suggesting they could be a site where anabolic stimuli convene to regulate mTORC1 kinase activity. Indeed, the forced targeting of mTORC1 to focal adhesion complexes elevates mTORC1 activity irrespective of lysosomal positioning (22). In skeletal muscle paxillin, a commonly used marker of focal adhesion complexes, is observed to be colocalized with the microvasculature (23), a region which mTORC1 has been seen to translocate to in response to anabolic stimuli (17) and where amino acid transporters (e.g. L-type amino acid transport) reside (24). Therefore, focal adhesion complexes may also contribute to mTORC1 activation in human skeletal muscle, however, a direct association between the two has yet to be observed.

The primary aim of the current investigation was to establish a method to visualize mTORC1 activity in human skeletal muscle and explore the effects of anabolic stimuli on the localization of mTORC1 activity. In addition, we also aimed to determine if mTORC1 activity occurred in the vicinity of focal adhesion complexes and if mTORC1 activation was regulated in a fiber type-specific manner. To achieve this, we utilized p-RPS6^Ser240/244^ as a marker of mTORC1 activity as it is an mTORC1-specific event (25, 26) has regularly been shown to be rapamycin-sensitive in various models (5, 25, 27, 28). We hypothesized that mTORC1 activity would occur predominantly in the periphery of muscle fibers, the location to which mTORC1 is commonly observed to translocate. We also hypothesized that mTORC1 activity would be enriched in focal adhesion complexes and would occur to a similar extent in oxidative and glycolytic fibers.

## Materials and Methods

### Subjects and Ethical Approval

Fourteen young, healthy, recreationally active males (age; 23±4yrs, weight; 77.5±17.6kg, body fat (BF); 12.9±4.3%) volunteered to participate in the current study. All participants provided written informed consent after being informed of the procedures and risks associated with the study. Each participant completed a physical activity readiness questionnaire prior to enrollment to ensure they were healthy and able to complete resistance exercise. Exclusion criteria were: i) tobacco and/or illicit anabolic drug use (e.g. testosterone, growth hormones); ii) veganism or nut/dairy allergies and; iii) injuries preventing participation in weight lifting/resistance exercise. Participants were randomized to either consume a protein-carbohydrate beverage at rest (FED, n=7, 23±5yrs, 76.8±3.6kg, 13.6±3.8%BF), or following a bout of whole-body resistance exercise (EXFED, n=7, 22±2yrs, 78.1±3.6kg, 12.2±4.9%BF). All procedures were approved by the research ethics board at the University of Toronto, Canada (Protocol #00036752) and conformed to the Declaration of Helsinki (revised 2013).

### Preliminary assessments

Participants attended the laboratory 3-7 days prior to experimental trial days for body composition and maximal strength testing. Following an overnight (⁓10h) fast, and prior to water consumption, participants height and weight were measured with a stadiometer and calibrated scales respectively before body composition was assessed via air displacement plethysmography (BOD−POD, COSMED USA Inc., Chicago, IL, USA). Participants were then provided with a light snack and underwent maximal strength training for the following exercises: i) dumbbell chest press, ii) dumbbell row, iii) leg extension, and; iv) leg press. This testing consisted of a warm-up set at a self-selected load, followed by progressive load increments until participants could no longer complete one repetition with correct form. The final load at which participants successfully completed a repetition was recorded as the 1 repetition maximum (1RM) and was used to calculate loads to be used during the experimental trial.

### Experimental trial

On the day of the experimental trial, participants attended the laboratory following an overnight fast (⁓10h) and having refrained from strenuous exercise and alcohol consumption in the previous 48 hours. Participants rested in a supine position upon arrival for 10min after which a skeletal muscle biopsy sample (PRE) was obtained from the *vastus lateralis*, of a randomly selected leg, using the modified Bergström technique (29) under local anesthesia (2% lidocaine). Participants then undertook their assigned intervention, either consumption of a protein-carbohydrate beverage alone or following a whole-body resistance exercise bout. The protein-carbohydrate beverage contained crystalline amino acids in a formulation modeled on the composition of egg protein (30) and artificially flavored maltodextrin (Tang, Kraft Foods Inc. Chicago, Illinois, United States) providing 0.25 g/kg protein and 0.75g/kg carbohydrate. The whole-body resistance exercise bout consisted of 4 sets of each exercise (chest press, dumbbell row, leg press & leg extension) at 75%1RM completed to volitional failure (⁓8-12 repetitions) with a 2min rest interval between each set/exercise. Immediately following exercise cessation, EXFED participants consumed the protein-carbohydrate beverage. After consumption of the beverage participants remained supine for the following 300min with subsequent skeletal muscle biopsy samples obtained at 120 and 300min from separate incisions. Muscle biopsy samples were freed from any visible blood, adipose, and connective tissue and placed in optimal cutting temperature compound (VWR International, Mississauga, ON, Canada) and frozen in liquid nitrogen-cooled isopentane before storage at −80°C for immunofluorescence analysis. A separate piece of the muscle biopsy sample immediately frozen in liquid nitrogen and stored at −80°C until subsequent immunoblot analysis.

### Immunoblotting

Immunoblotting was completed as described previously (19). Briefly, a small piece of skeletal muscle tissue (20mg) was homogenized by mechanical pulverization in radioimmunoprecipitation assay (RIPA) buffer (65 mM Tris-base, 150 mM NaCl, 1% NP-40, 0.5% sodium deoxycholate, 0.1% sodium dodecyl sulfate) with added protease and phosphatase inhibitors (Roche Applied Science, Mannheim, GER). Myofibrillar proteins were pelleted by centrifugation at 700g for 5min and the protein concentration of the sarcoplasmic fraction (supernatant) was determined via a bicinchoninic acid assay (Thermo Fisher Scientific, Rockford, IL). Samples were then diluted to equal concentrations in 1X Laemmli sample buffer and denatured at 95℃ for 5min. Equal amounts of protein were then separated by SDS-PAGE on 4-20% polyacrylamide gels (Bio-Rad Laboratories, Richmond, VA, USA) at 200V for 40min, and proteins were transferred onto nitrocellulose membranes at 100V for 1h. Membranes were then stained with Ponceau S to confirm equal loading and blocked in 5% (wt/vol) bovine serum albumin in Tris‐buffered saline with 0.1% Tween 20 (TBST) for 1h at room temperature (RT). Following this, membranes were incubated in primary antibody (rpS6^Ser240/244^ #5364, tRPS6 #2217, Cell Signaling, Danvers, MA.), diluted 1:1000 in TBST, overnight at 4℃. Membranes were then washed in TBST and incubated in horseradish peroxidase-conjugated secondary antibody (1:10000 in TBST, Cell Signaling) for 1h at RT. Bands were detected with chemiluminescent substrate (WBKLS0500, Millipore, Etobicoke, ON, Canada) and visualized using a Fluorochem E Imaging system (Protein Simple; Alpha Innotech, Santa Clara, CA). Bands were quantified using Protein Simple AlphaView SA software and normalized to Ponceau S and a gel control (identical generic sample run on every gel).

### Immunofluorescence

Serial skeletal muscle cross sections (7μm) were sectioned onto room temperature uncoated glass slides (ThermoFisher SuperFrost+, Fisher Scientific, Rockford, IL) and air-dried to remove excess water. For p-RPS6^Ser240/244^ staining, sections were fixed in 4% paraformaldehyde for 10min at 4℃ to create covalent cross links. Sections were then washed in 1xPBST (PBS supplemented with 0.2% Tween20) and incubated in a blocking solution consisting of 2% bovine serum albumin (BSA), 5% fetal bovine serum, 5% normal goat serum (NGS), 0.2% Triton X-100 and 0.1% sodium azide for 90min at RT. Following this, sections were washed again in 1xPBST for 5min and incubated in primary antibody, diluted in 1%BSA, overnight at 4℃ (antibodies and dilutions are listed in Table 1.). The next day, following further 1xPBST washes (3×5min), sections were incubated in corresponding secondary antibodies (1:300 dilution in 1%BSA in 1xPBS, detail in Table 1) for 2h at RT. Sections were then either incubated in wheat germ agglutinin (WGA, 1:20 in 1%BSA), to mark the cell membrane (Myosin heavy chain 1-RPS6^Ser240/244^ stain only), for 30min or DAPI (1:10000 in 1xPBS), to mark nuclei, for 10min and then dried and covered DAKO fluorescent mounting medium (Agilent, Santa Clara, CA) and a coverslip prior to imaging. To confirm specificity of the p-RPS6^Ser240/244^ antibody to phosphorylated forms of the protein, a lambda protein phosphatase (LPP) assay was completed (n=2, 120 & 300 time points only for each condition). Here, prior to fixation, tissue sections were incubated in a solution of LPP, NEBuffer and Manganese Chloride (MnCl2) as per manufacturer’s instructions (P0753, Cell Signaling) for 2h. Following this, p-RPS6^Ser240/244^ immunofluorescence staining was completed as described above. A duplicate tissue section on the same slide was stained as normal (i.e. no LPP incubation) to act as a comparative control. To ensure fluorescence bleed through was not affecting results during fiber type staining, a subset of samples (n=2 per condition) were stained via serial staining and co-staining methods simultaneously. For serial staining, one section was stained for Myosin heavy chain 1 (MHC1) and WGA whilst a second identical section on the same slide was stained for p-RPS6^Ser240/244^. The first section was then used to identify fibers on the second section and staining intensity measured, and compared to a co-stain on the same slide, in order to assess if the presence of MHC1 on the same section as p-RPS6^Ser240/244^ affected outcomes.

**Table 1.** Summary of Antibodies Used.

| Primary Antibody | Source | Dilution | Secondary Antibody | Dilution |
| --- | --- | --- | --- | --- |
| Monoclonal anti-pRPS6<sup>Ser240/244</sup> antibody with rabbit antigen, isotype IgG | Cell Signaling Technology #5364 | 1:50 | Goat anti-rabbit IgG(H+L) Alexa@488 | 1:300 |
| Monoclonal anti-Dystrophin antibody with mouse antigen, isotype IgG2a | DSHB, MANDYS1 3B7 | 1:200 | Goat anti-mouse IgG(H+L) Alexa@594 | 1:300 |
| Monoclonal anti-MHC1 antibody with mouse antigen, isotype IgG γ1 kappa | DSHB, A4.951 | Neat | Goat anti-mouse IgG γ1 kappa Alexa@594 | 1:300 |
| Monoclonal anti-Paxillin antibody (Clone 349) with mouse antigen, isotype IgG γ1 kappa | Millipore, MAB3060 | 1:500 | Goat anti-mouse IgG γ1 kappa Alexa@594 | 1:300 |
| Monoclonal anti- mTOR antibody with mouse antigen, isotype IgG γ1 kappa | Millipore, 05-1592 | 1:200 | Goat anti-mouse IgG γ1 kappa Alexa@594 | 1:300 |
| DAPI (4',6-Diamidino-2-Phenylindole, Dihydrochloride) | Invitrogen, D1306 | 1:10,000 | N/A | N/A |
| Wheat Germ Agglutinin-350 | Invitrogen, W11263 | 1:20 | Alexa Fluor® 350 Conjugated | N/A |

For mTOR staining, sections were fixed in an acetone-ethanol solution (3:1 ratio) for 5min and blocked with 5%NGS for 1h at RT. Samples were then incubated in primary antibodies (Table 1) for 2h at RT, washed in 1xPBST (3×5min), and exposed to relevant secondary antibodies (1:300 dilution in 1xPBS, Table 1) for 1h at RT. WGA was then used to mark the sarcolemma (1:20 in 1% BSA) for 30min at RT and slides were then covered with DAKO fluorescent mounting medium and a coverslip prior to imaging.

### Image capture and analysis

Image capture was undertaken on an EVOS FL Auto Cell imaging microscope (Thermo Fisher Scientific, Waltham, MA) at 40× 0.75NA magnification, using automated image capture and stitching functions. All sections from a single participant were stained and imaged on a single slide and all image capturing parameters were kept constant for all images including exposure time, gain, and light intensity. As such, images containing approximately 100 fibers were captured for each time point, for each participant for subsequent analysis.

Image processing and quantification was all completed using Image J (Fiji plugin, v. 1.5, National Institutes of Health, USA). For total staining intensity analysis, mean pixel intensity was measured across the entire section. For peripheral staining intensity, mean pixel intensity in the outer 5.5μm of each skeletal muscle fiber was assessed, with the remaining region of the fiber being recorded as the ‘central’ region. This region was chosen as it is the region within which mTORC1 is observed in following anabolic stimuli (16, 17, 20). Peripheral-central ratio was calculated as mean pixel intensity in the peripheral region divided by the mean pixel intensity in the central region. To determine fiber-type specific staining of p-RPS6^Ser240/244^, type I fibers were recorded as those with positive myosin heavy chain 1 staining whereas all ‘negative’ fibers were recorded as type II fibers. When assessing staining intensity within paxillin-positive regions, thresholding was used to determine paxillin positive regions of the cell and regions of interest created in these areas. These areas were then transferred to p-RPS6^Ser240/244^ channel images and mean pixel intensity was measured. Thresholding levels for paxillin were kept identical across all images for each participant. Finally, colocalization analysis was conducted by quantifying the Pearson’s correlation coefficient where each individual pixel’s intensity in each channel was plotted and the corresponding r value calculated. Therefore, increases in colocalization between two targets at a given time point would result in *r* values moving closer to 1.

### Statistical analysis

All statistical tests were conducted in SPSS statistics version 24 for Windows (IBM, Armonk, NY, USA), with significance set at P < 0.05. Two-way repeated measures analysis of variance (ANOVA) tests with one within-subject factor (time point) and one between-subject factor (condition) were conducted for all outcome measures other than fiber-type specific RPS6^Ser240/244^ staining intensities. For this measure, a three-way repeated measures ANOVA was conducted, with two within-subject (time and fiber type) and one between-subject (condition) factor. If a statistical test failed Mauchly’s test of sphericity, a Greenhouse-Geisser correction was used. If assumptions of normality (Shapiro-Wilk test) were violated, logarithmic transformed values were used. If a significant main or interaction effect was observed, *post hoc* t-tests were conducted, in Microsoft Excel, with a Holm-Bonferroni correction for multiple comparisons. To tests associations between outcome variables, pearson’s r correlation coefficients were calculated in GraphPad version 8.00 for Windows (GraphPad Software, San Diego, CA). All data is presented as Mean±SD unless stated otherwise.

## Results

### Confirmation of p-RPS6^Ser240/244^ stain

The p-RPS6^Ser240/244^ immunofluorescent staining protocol was confirmed by the use of a LPP assay. Incubation of sections with LPP significantly reduced p-RPS6^Ser240/244^ staining intensity confirming the specificity of the antibody to phosphorylated RPS6 (p<0.001, Figure 1A). The stain was further confirmed via the omission of primary or secondary antibodies to determine non-specific secondary antibody staining and contribution of autoflourescence respectively. When either antibody was omitted, staining intensity was no longer apparent in muscle cross-sections imaged using identical parameters to a control stain run on the same sample (Figure 1B & 1C).

**Figure 1.**
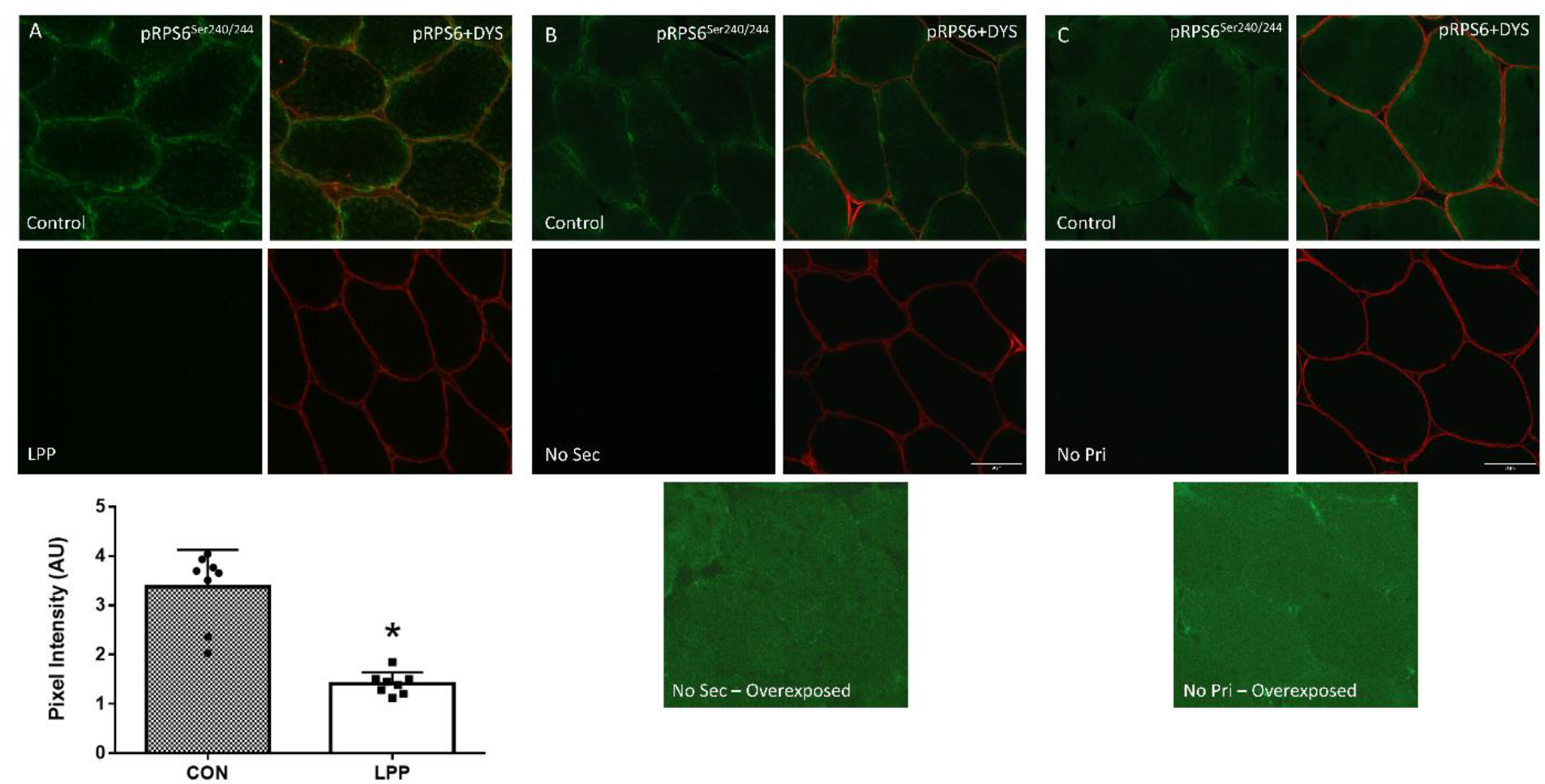
Validation of the p-RPS6^Ser240/244^ immunofluorescent stain via Lambda protein phosphatase assay (LPP) (A) and secondary antibody (B) and primary antibody (C) omission stains. *Significantly different from CON (p<0.001).

### RPS6^Ser240/244^ phosphorylation assessed by immunoblot

A time x condition interaction effect was observed for p-RPS6^Ser240/244^ (p=0.039). RPS6^Ser240/244^ phosphorylation was elevated at 120min following both FED and EXFED (5.3±3.1 and 21.3±15.6fold, respectively; p<0.05), before returning to PRE at 300min (Figure 2A). The extent of RPS6^Ser240/244^ phosphorylation in EXFED was also greater than that in FED at both 120 and 300min time points (21.3±15.6 vs. 5.3±3.1fold and 4.4±6.4 vs. 0.97±0.79 fold, respectively; p<0.05, Figure 2A). A condition effect was found for total RPS6 protein content (p=0.011) where, irrespective of time point, FED individuals had greater expression of total RPS6 (Figure 2B). No time (p=0.75) or interaction (p=0.403) effect was apparent. When p-RPS6^Ser240/244^ was expressed in relation to total RPS6, a time x condition effect was observed (p=0.001). p-RPS6^Ser240/244^/t-RPS6 was elevated at 120min following both FED and EXFED (4.5±2.0 and 31.1±15.5fold, respectively; p<0.01), with the response in EXFED greater than FED at the time point (p<0.001, Figure 2C). RPS6^Ser240/244^/t-RPS6 then returned to baseline levels in FED at 300 (0.95±0.7fold, p>0.05) but remained above baseline and FED at the same time point in EXFED (8.8±11.5fold, p=0.047, Figure 2C).

**Figure 2.**
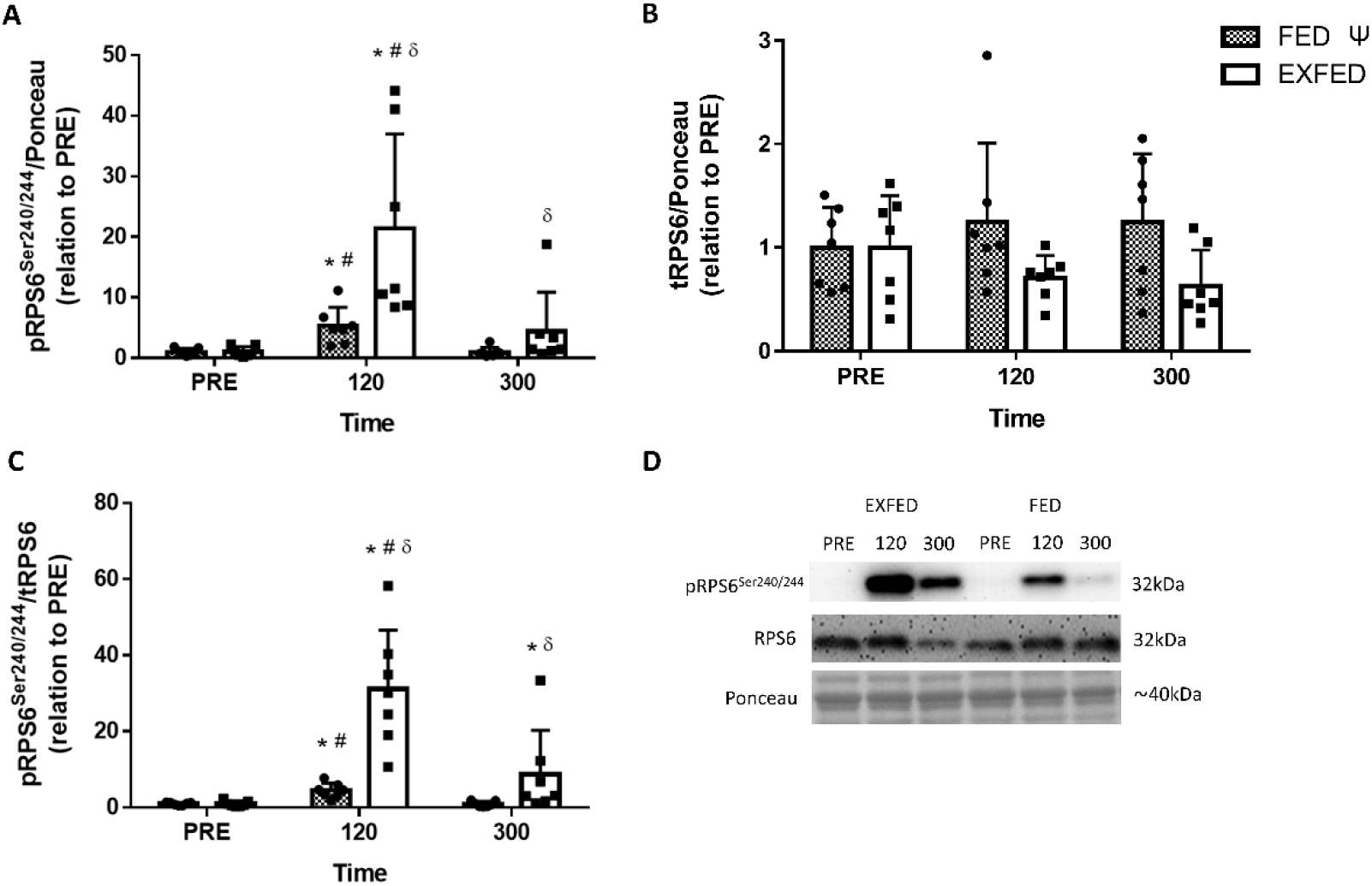
The effect of protein-carbohydrate beverage ingestion at rest (FED) or following resistance exercise (EXFED) on RPS6^Ser240/244^ phosphorylation (A), total RPS6 protein content (B) and RPS6^Ser240/244^ phosphorylation in relation to total protein content (C). Representative immunoblot images are shown in panel D. *significantly different from PRE, ^#^significantly different from 300, ^δ^significant difference between conditions, ^Ψ^significant condition effect (p<0.05). Data presented as Mean±SD, n=7 per condition.

### p-RPS6^Ser240/244^ total staining intensity and correlation to immunoblots

Total RPS6^Ser240/244^ phosphorylation measured by immunofluorescence staining displayed a time x condition interaction effect (p=0.027). p-RPS6^Ser240/244^ staining intensity was elevated in both FED and EXFED at 120min (50±22% and 103±40%, respectively, p<0.05, Figure 3A), with this increase being greater in EXFED (p=0.02). In FED, p-RPS6^240/244^ staining intensity returned to PRE at 300min, however remained above PRE levels in EXFED (25±18%, p=0.04, Figure 3A). In addition, at 300min a trend suggesting EXFED induced greater p-RPS6^Ser240/244^ staining intensity compared to FED (25±18% vs 8±13%, p=0.06). When comparing p-RPS6^Ser240/244^ measured by immunofluorescent staining and immunoblotting, a strong positive association was apparent (r=0.76, p<0.001. Figure 3B).

**Figure 3.**
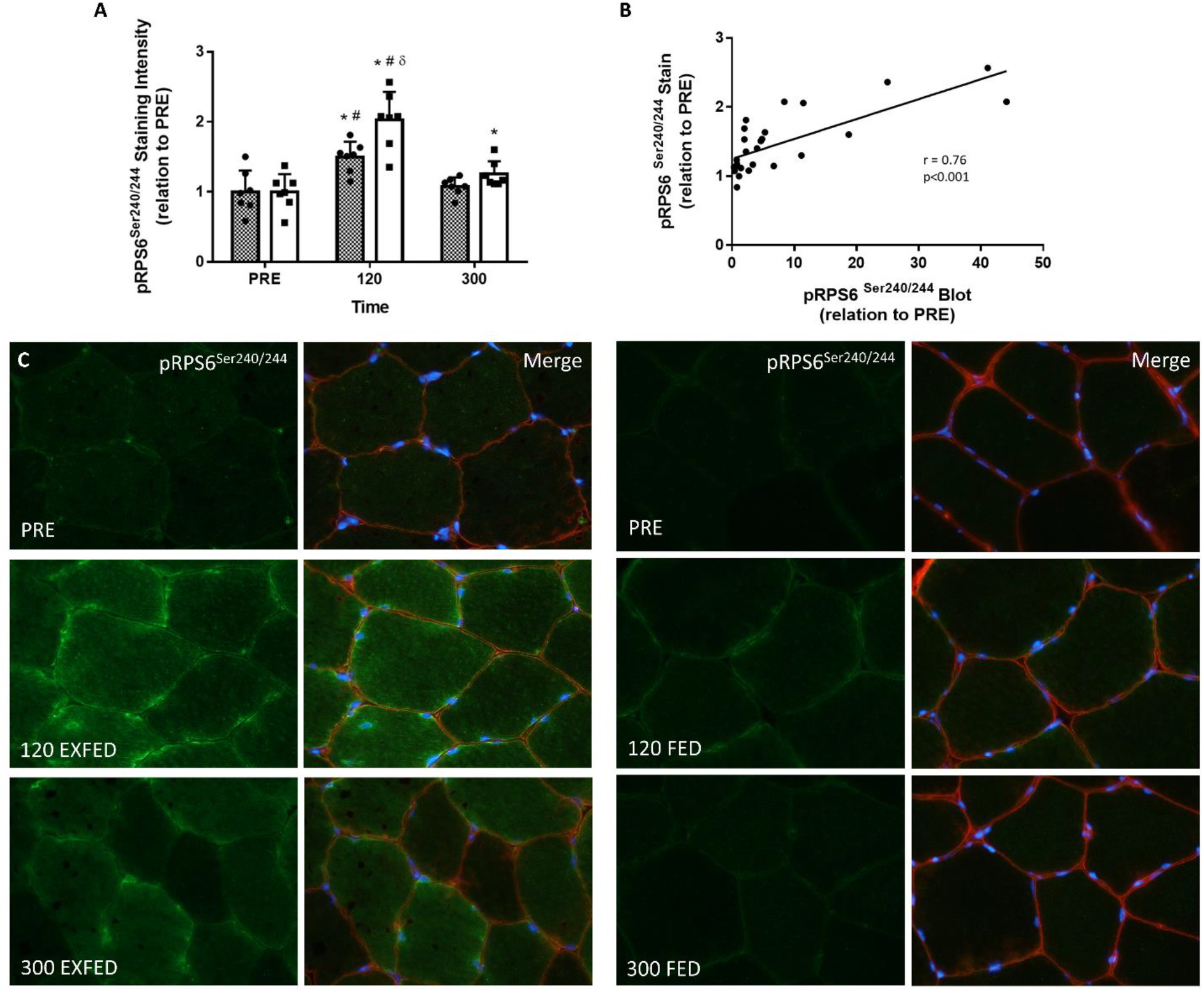
The effect of protein-carbohydrate beverage ingestion at rest (FED) or following resistance exercise (EXFED) on RPS6^Ser240/244^ phosphorylation measured by immunofluorescence microscopy (A). Pearson’s correlation coefficient analysis between immunoblot and immunofluorescent staining is also shown (B). Representative images of the p-RPS6^Ser240/244^ stains are shown in (C), with p-RPS6^Ser240/244^ alone and merged with dystrophin and DAPI provided. *significantly different from PRE, ^#^significantly different from 300, ^δ^significant difference between conditions at this timepoint (p<0.05). Data presented as Mean±SD, n=7 per condition.

### Region-specific RPS6^Ser240/244^ phosphorylation

When examining RPS6^Ser240/244^ phosphorylation in central regions of fibers a time x condition interaction effect was observed (p=0.035). p-RPS6^Ser240/244^ was elevated at 120min only in EXFED (92±40%, p<0.01) and was greater compared to FED (44±19%) at this time point (p=0.026, Figure 4A). There was also a trend for central RPS6^Ser240/244^ phosphorylation in FED to be greater than PRE at 120min (p=0.054, Figure 4A). No differences from PRE, or between conditions, was apparent at 300min (p>0.05). A time x condition interaction was also observed for peripheral p-RPS6^Ser240/244^ staining intensity (p=0.001). Peripheral p-RPS6^Ser240/244^ increased in both FED and EXFED at 120min (54±23% & 137±48% respectively, p<0.05), but to a greater extent in EXFED (p=0.003, Figure 4B). At 300min, peripheral RPS6^Ser240/244^ phosphorylation remained above PRE in EXFED only (34±26%) and was greater than FED (7±13%) at this time point (p=0.033, Figure 4B). As RPS6^Ser240/244^ phosphorylation was elevated in both central and peripheral regions, particularly in EXFED, peripheral-central ratio was calculated to determine if mTORC1 activity occurred to a greater extent in either region. A time x condition interaction was also apparent for this variable (p<0.001), whereby p-RPS6^Ser240/244^ peripheral-central ratio increased, above PRE, in both FED and EXFED at 120min (1.20±0.05 to 1.28±0.06AU in FED, 1.17±0.03 to 1.44±0.08AU in EXFED, p<0.05, Figure 4C). Peripheral-central ratio of RPS6^Ser240/244^ phosphorylation remained above PRE at 300min only in EXFED (1.26±0.08AU, p=0.021), and peripheral-central ratios at both 120 and 300min were greater in EXFED compared to FED (p<0.05, Figure 4C).

**Figure 4.**
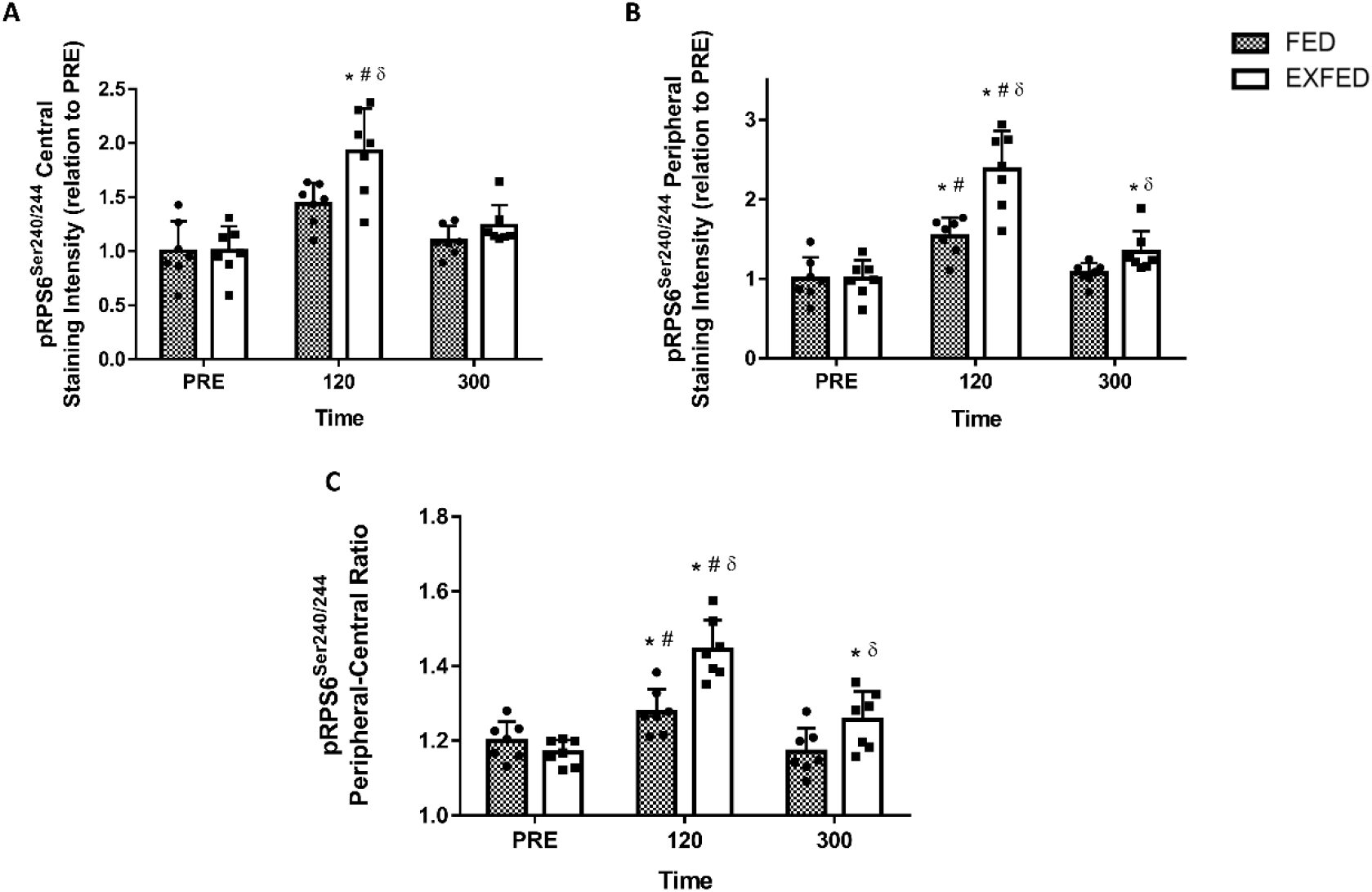
The effect of protein-carbohydrate beverage ingestion at rest (FED) or following resistance exercise (EXFED) on RPS6^Ser240/244^ phosphorylation in central (A) and peripheral (B) regions of fibers. Peripheral-central ratio of p-RPS6^Ser240/244^ staining in each condition is also shown (C). *significantly different from PRE, ^#^significantly different from 300, ^δ^significant difference between conditions (p<0.05). Data presented as Mean±SD, n=7 per condition.

### Fiber type specific RPS6^Ser240/244^ phosphorylation

When presented as total p-RPS6^Ser240/244^ pixel intensity in type I and type II fibers, only time (p<0.001) and fiber type effects were observed (p<0.001). Accordingly, p-RPS6^Ser240/244^ staining intensity was elevated at 120min compared to PRE and 300min (both p<0.01) irrespective of condition or fiber type (Figure 5A). Furthermore, irrespective of time or condition, p-RPS6^Ser240/244^ staining intensity was greater in type I fibers (33±24% overall, Figure 5A). When presented in relation to PRE staining intensities, only a significant time effect was apparent (p<0.001) showing that, irrespective of condition or fiber type, RPS6^Ser240/244^ phosphorylation was elevated at 120min compared to PRE and 300min (both p<0.01, Figure 5B). This suggests that after correcting for basal differences, there are no fiber type differences in RPS6^Ser240/244^ phosphorylation in response to FED or EXFED. To confirm these differences were not a result of fluorescence bleed through between channels, fiber type specific RPS6^Ser240/244^ staining was assessed by serial section staining and direct co-staining in n=2 subjects per condition and compared. Here, a strong positive correlation was found between methods of staining (r=0.924, p<0.001, data not shown) suggesting differences were not a result of fluorescence bleed through.

**Figure 5.**
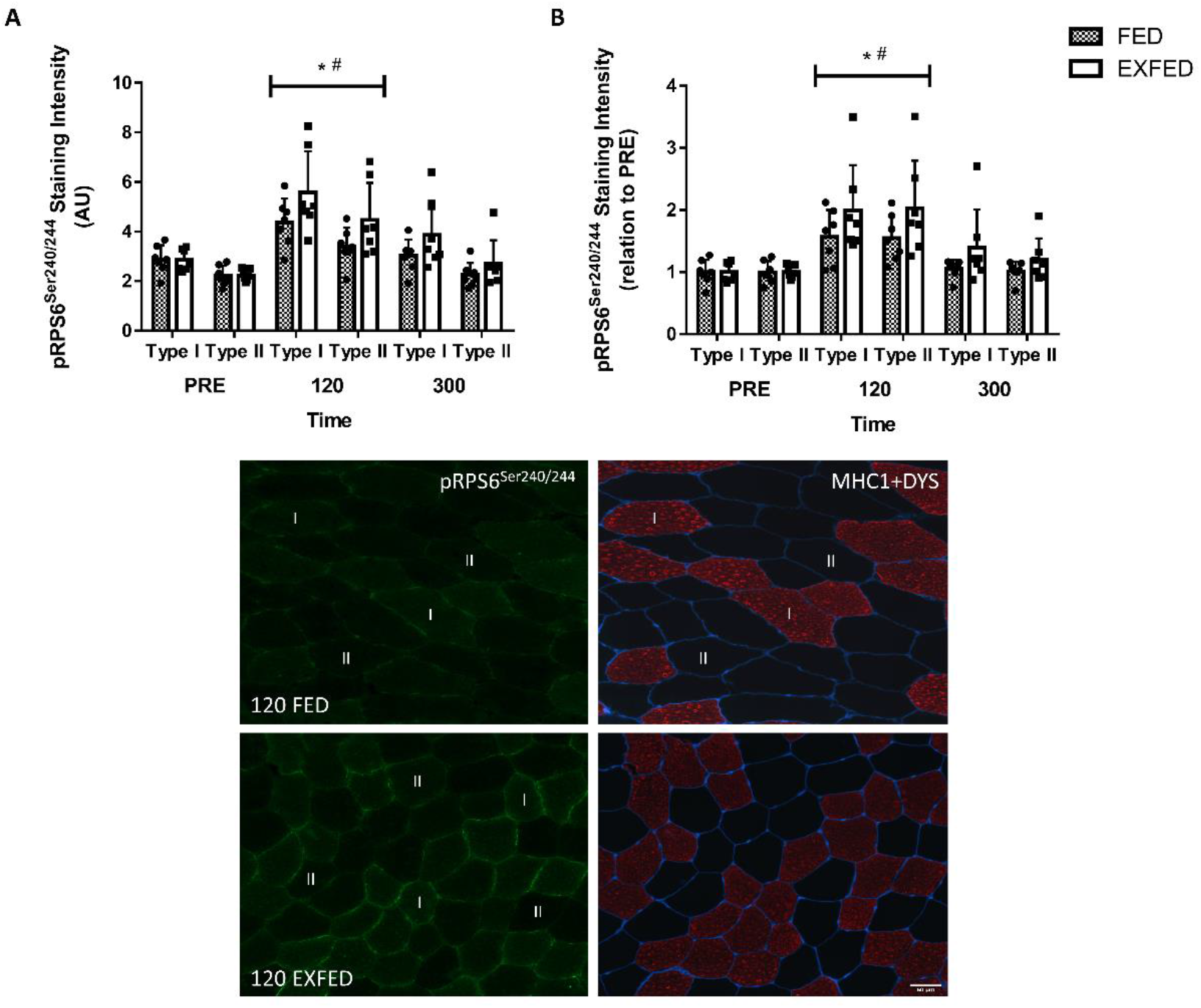
The effect of protein-carbohydrate beverage ingestion at rest (FED) or following resistance exercise (EXFED) on fiber type-specific RPS6^Ser240/244^ phosphorylation presented as arbitrary units (A) or relative to PRE (B). Representative images of the 120 timepoint are shown in (C), with p-RPS6^Ser240/244^ alone (green) and Myosin Heavy Chain 1 + dystrophin (red + blue) provided. I indicates type I fibers and II indicates type II fibers on representative images. *significantly different from PRE, ^#^significantly different from 300 (p<0.05). Data presented as Mean±SD, n=7 per condition.

### RPS6^Ser240/244^ phosphorylation in proximity to focal adhesion complexes

p-RPS6^Ser240/244^-Paxillin colocalization analysis revealed a significant time (p=0.036) but not condition (p=0.787) or time x condition interaction (p=0.66) effect. Irrespective of condition, p-RPS6^Ser240/244^-Paxillin colocalization was greater at 120min compared to 300min (p=0.046, Figure 6A). Furthermore, there was also a trend toward RPS6^Ser240/244^-Paxillin colocalization being greater at 120min, compared to PRE, irrespective of condition (p=0.051, Figure 6A). p-RPS6^Ser240/244^ association with focal adhesion complexes was also assessed via the measurement of p-RPS6^Ser240/244^ staining intensity with paxillin ‘positive’ regions. A significant effect of time (p=0.001) showed significantly greater RPS6^Ser240/244^ phosphorylation within these regions at 120min (FED – 45±34%, EXFED – 91±54%) compared to both PRE and 300min (both p<0.01, Figure 6B). In addition, a trend toward a condition effect (p<0.074) was found suggesting that, irrespective of time point, p-RPS6^Ser240/244^ staining intensity in paxillin ‘positive’ regions was greater in EXFED compared to FED (Figure 6B).

**Figure 6.**
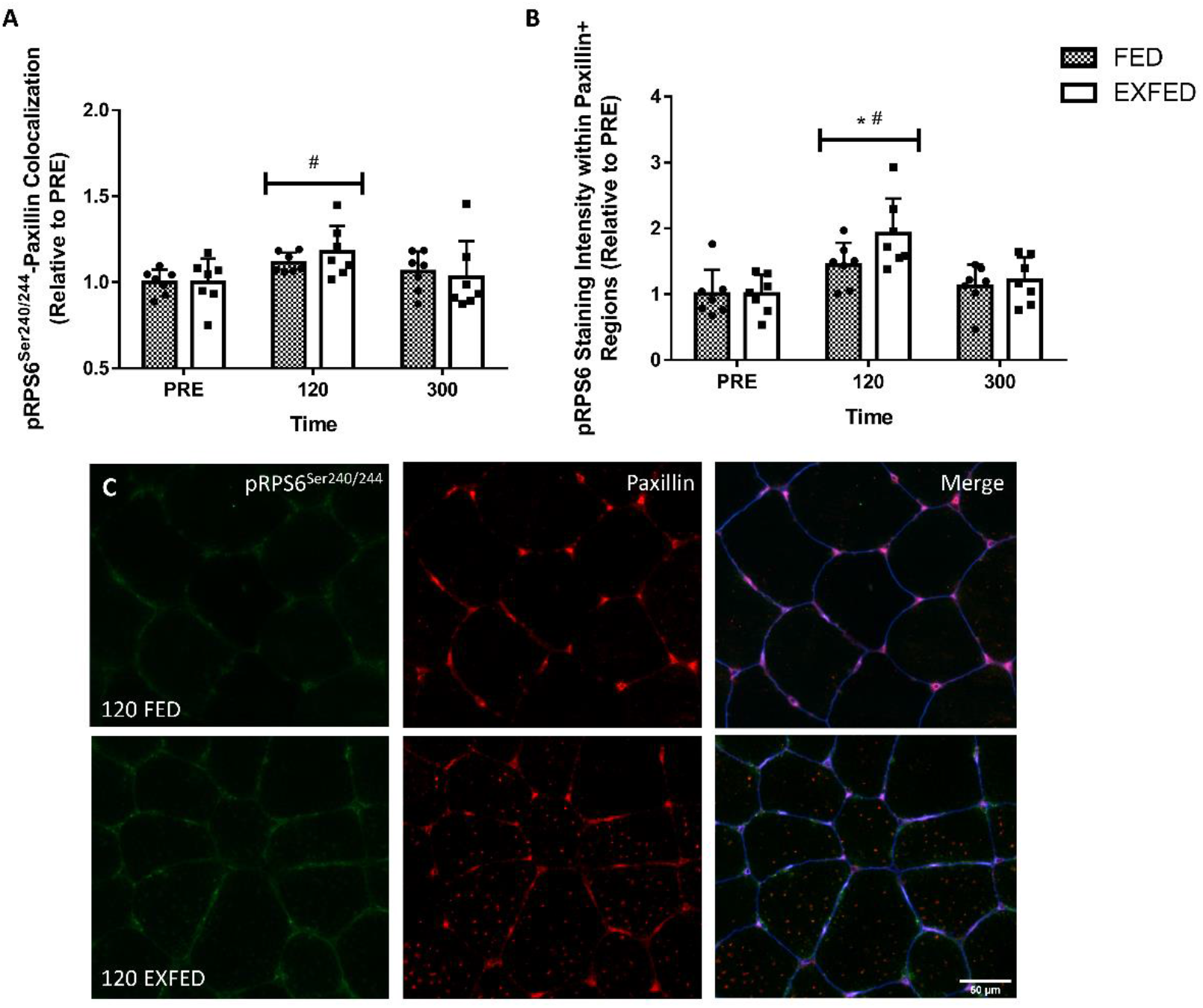
The effect of protein-carbohydrate beverage ingestion at rest (FED) or following resistance exercise (EXFED) on p-RPS6^Ser240/244^ association with focal adhesion complexes assessed as p-RPS6^Ser240/244^-Paxillin colocalization (A) and p-RPS6^Ser240/244^ staining intensity with paxillin-positive regions (B). Representative images of the 120 timepoint are shown in (C), with p-RPS6^Ser240/244^ alone (green), paxillin alone (red) and p-RPS6^Ser240/244^, paxillin and WGA overlayed (Merge) provided. *significantly different from PRE, ^#^significantly different from 300 (p<0.05). Data presented as Mean±SD, n=7 per condition.

### Confirmation of mTORC1 translocation to cell periphery

In order to confirm mTORC1 translocation, both dystrophin and WGA were used as markers of the cell periphery. A condition effect was observed for mTOR-WGA colocalization (p=0.004) displaying that, irrespective of time point, this measure was greater in EXFED compared to FED (Figure 7A). A similar effect of condition was also noted for mTOR-dystrophin colocalization (p=0.02) where it was greater in EXFED compared to FED, irrespective of time point (Figure 7B). mTOR-WGA and mTOR-dystrophin colocalization were then compared to assess agreeability between the two measures. A strong, positive correlation between the two measures was observed (r=0.77, p<0.001, Figure 7C), suggesting both WGA and dystrophin are reliable markers of the cell periphery to assess mTORC1 translocation. Finally, mTOR peripheral-central ratio was assessed as a further measure of mTOR peripheral content. A time x condition interaction effect was observed (p=0.015), with mTOR peripheral-central ratio increasing above PRE at 120 and 300 in EXFED (PRE – 1.16±0.03AU, 120 – 1.24±0.05AU, 300 – 1.23±0.04AU, p<0.05, Figure 7D). In FED, the only difference apparent was a greater mTOR peripheral-central ratio at 120min compared to 300min (1.26±0.06AU vs. 1.20±0.07, p=0.016, Figure 7 D). No differences between FED and EXFED at any time point was apparent (p>0.05).

**Figure 7.**
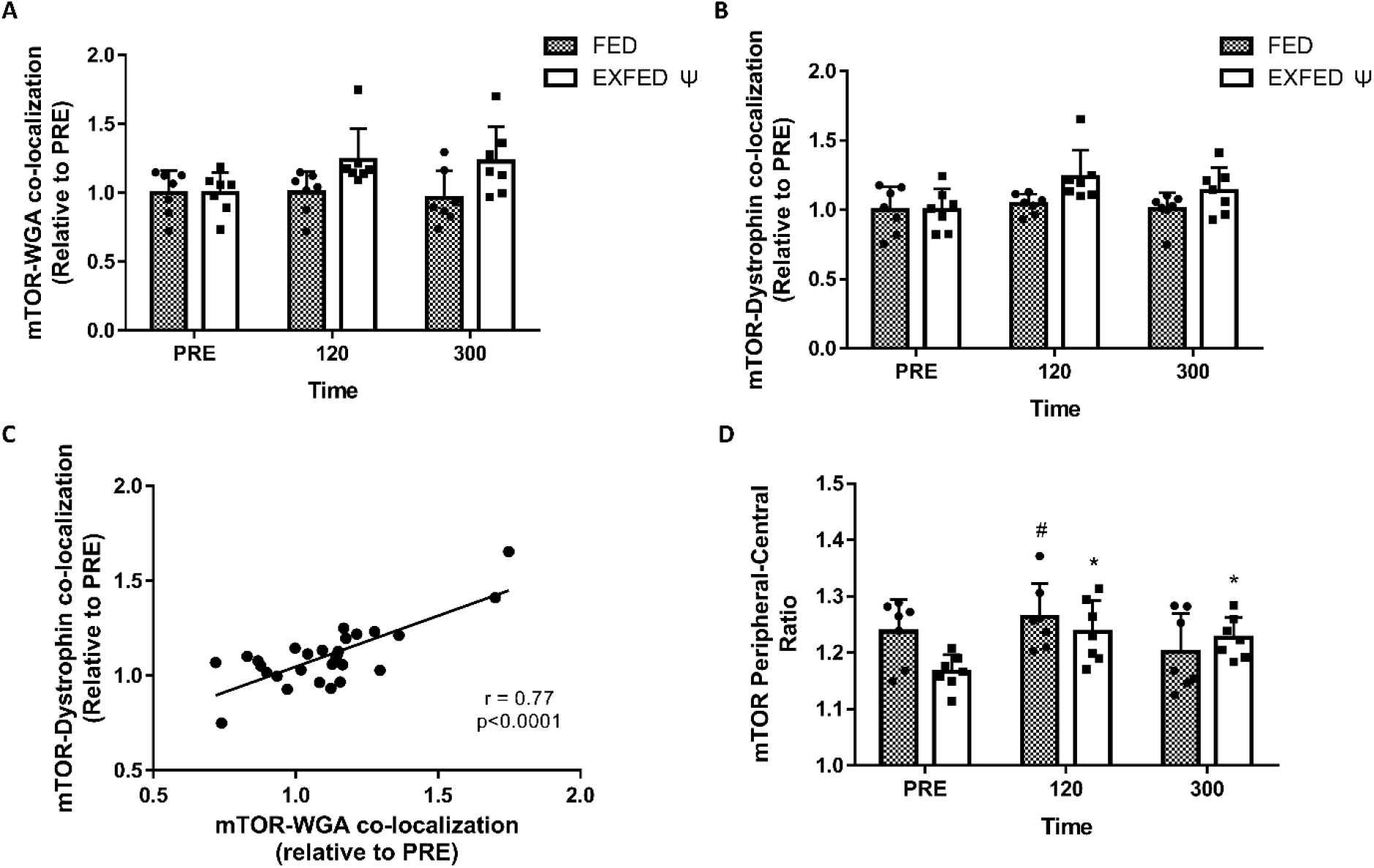
The effect of protein-carbohydrate beverage ingestion at rest (FED) or following resistance exercise (EXFED) on mTOR translocation assessed by mTOR-WGA colocalization (A), mTOR-Dystrophin colocalization (B) and peripheral-central ratio of mTOR staining (D). Pearson’s correlation coefficient analysis between the mTOR-WGA and mTOR-Dystrophin colocalization is also shown (C). *significantly different from PRE, ^#^significantly different from 300, ^δ^significant effect of condition (p<0.05). Data presented as Mean±SD, n=7 per condition.

## Discussion

Herein, we provide a novel method for the visualization of a marker of mTORC1 activity in human skeletal muscle. Using this method, we show, for the first time, that mTORC1 activity occurs predominantly in the periphery of human skeletal muscle fibers following anabolic stimuli and to a greater extent following EXFED compared to FED. In addition, when accounting for basal differences, anabolic stimuli do not regulate RPS6^Ser240/244^ phosphorylation in a fiber type-specific manner over the time period measured. In agreement with recent *in vitro* data (22), p-RPS6^Ser240/244^ was observed to localize with focal adhesion complexes, purported to be central regulators of anabolic signal transduction.

RPS6^Ser240/244^ phosphorylation, when measured by immunoblot, is elevated upwards of 5-fold following anabolic stimuli (5, 19, 31, 32), with these elevations often greater than other commonly assessed mTORC1-regulated sites such as p-S6K1^Thr389^ and p-4EBP1^Thr37/46^ (33). Furthermore, RPS6^Ser240/244^ phosphorylation has consistently been shown to be rapamycin-sensitive *in vitro* (25, 26) and in rodent and human skeletal muscle (5, 27, 28). In raptor-KO mice, which exhibit impaired mTORC1 complex formation and kinase activity, RPS6^Ser240/244^ phosphorylation following maximal intensity contractions was reduced by approximately 83% (34). Importantly, in raptor-KO animals, approximately 10% of raptor protein remained, which may account for the absence of fully ablated RPS6^Ser240/244^ phosphorylation. As such, it is apparent that RPS6^Ser240/244^ phosphorylation is almost entirely mTORC1-dependent. Another commonly measured phosphorylation site on RPS6 (serine 235/236) is not rapamycin-sensitive in all models (25–27), most likely due to the ability of p90 ribosomal S6 kinase (p90RSK) to phosphorylate this site in addition to S6K1 (26). Collectively, these data show that RPS6^Ser240/244^ is a mTORC1-specific phosphorylation event that is robustly increased following anabolic stimuli in skeletal muscle and thus an ideal candidate to translate from immunoblot to immunofluorescent staining methods. The p-RPS6^Ser240/244^ stain developed here was first confirmed using a LPP assay which removes all phosphate from proteins within a tissue section. In the presence of LPP, p-RPS6^Ser240/244^ staining intensity was greatly reduced to levels similar to those observed on unstained skeletal muscle sections at the FITC wavelength (488nm). These data confirm that the antibody utilized for this stain is specific to phosphorylated forms of protein. We then further confirmed the staining pattern via primary and secondary antibody omission stains which confirmed that neither autofluorescence nor non-specific secondary antibody staining contributed extensively to the staining pattern observed. Although these control measures can definitively confirm staining specificity to and epitope on RPS6 (35), we interpret the strong, positive correlation found between p-RPS6^Ser240/244^ measured by immunoblot or immunofluorescent staining (r=0.76, Figure 2C) as supportive of our stain being a reliable readout of this phosphorylation event and therefore mTORC1 activity.

Using this newly developed immunofluorescent staining protocol, we observed that total RPS6^Ser240/244^ staining intensity was elevated in both FED and EXFED conditions at 120 min, however the increase in EXFED was greater. This finding is in agreement with a multitude of previous papers demonstrating that several readouts of mTORC1 activity are elevated by amino acid/protein ingestion but are greater and/or more prolonged after a bout of resistance exercise, highlighting mTORC1 activity is greater than with either anabolic stimulus alone (20, 36–38). We then aimed to determine the spatial localization of mTORC1 activity. Seminal *in vitro* research identified a mechanism of mTORC1 activation centering on the lysosome where, in response to amino acid and growth factor exposure, mTORC1 translocates to the lysosomal surface and is in close vicinity to direct activators such as Rheb and phosphatidic acid (8, 10, 39). However, we and others (16–18, 20, 21, 40) have observed a different mechanism where mTORC1 does not dissociate form the lysosomal membrane during fasting/nutrient deprivation and therefore mTORC1-lysosomal colocalization is not elevated in response to anabolic stimuli. Instead, following nutrient ingestion or mechanical loading, mTORC1-lysosome complexes translocate toward the periphery of the cell and colocalize with upstream activators (e.g. Akt and Rheb) and downstream substrates (e.g. translation initiation factors) (17, 18, 40). Although this new mechanism has been observed in several different investigations in human skeletal muscle (17, 20, 21, 40), it has yet to be confirmed if this translocation culminates in mTORC1 activation. Here, we show that RPS6^Ser240/244^ phosphorylation occurred in peripheral regions following both anabolic stimuli and in central regions of fibers following EXFED only at 120 min. Interestingly, the peripheral-central ratio was also elevated in both FED and EXFED suggesting mTORC1 activity increased to a greater extent in peripheral regions. This suggests that mTORC1 activation is occurring mainly in the regions where mTORC1 translocates to post-exercise/feeding, confirming it is most likely following/as a result of such translocation events. This has also been reported in *in vitro* investigations that the inhibition of lysosomal movement, via ablation of kinesin factors, impairs mTORC1 activation in response to nutrients (18). Importantly, the synergistic effect of EXFED was apparent for RPS6^Ser240/244^ peripheral-central ratio as this measure was greater in EXFED compared to FED at both 120 and 300 min timepoints. Although unexpected, an increase in central mTORC1 activity, in response to nutrients, has been observed *in vitro* (22). Intramyofibrillar ribosomes have also been identified in rat skeletal muscle, albeit at approximately 75-90% lower expression to subsarcolemmal, peripheral ribosomes (41) and, assuming similar ribosome localization in human muscle, it is therefore possible that mTORC1 activity also occurred in these regions following EXFED in order to stimulate myofibrillar protein turnover. Measures of mTOR translocation were also greater following this anabolic stimuli, confirming this event had also occurred in our current model, similar to our lab’s previous research (17, 20, 21, 40). Based on previous confirmatory data from our group (21) we take translocation of mTOR protein, in response to acute anabolic stimuli, as a measure of mTORC1 movement as we previously observed alterations only in Raptor cellular location in human skeletal muscle. These data therefore improve our current knowledge of mTORC1 activation in human skeletal muscle by confirming the notion that translocation of this kinase complex occurs prior to mTORC1 activation.

To our knowledge, this is the first study that has investigated the localization of mTORC1 activity in human skeletal muscle. However, several studies in rodent (42, 43) and human skeletal muscle (36, 44) have explored readouts of mTORC1 activity by immunofluorescent staining. In human skeletal muscle, protein-carbohydrate ingestion following resistance exercise elevated p-RPS6^Ser235/236^ staining intensity to a greater extent that carbohydrate ingestion alone post-exercise (36). Furthermore, a single night of wheel running (with 60% braking resistance) was able to significantly elevate RPS6^Ser235/236^ staining intensity in rodents (43). Importantly, although this phosphorylation event is not completely mTORC1-specific (25–27), the peripheral staining patterns observed in these studies were similar to that which we observed in the current study, although they were not quantified in previous work (36, 43). This staining pattern has also been observed in rodent skeletal muscle when assessing the same phosphorylation site as we probed here (42), however again regional-specific phosphorylation was not quantified. These data therefore add further credence to the notion that mTORC1 activity predominantly occurs in the periphery of skeletal muscle fibers. Additional evidence of this is also shown by the identification of the cell periphery in skeletal muscle as the main site of ribosomes (41), the organelles which drive protein synthesis, as well as the visualization of a strong puromycin (protein synthetic marker in SUnSET technique) signal close to the plasma membrane in rodent skeletal muscle fibers (45). The importance of the location of translation has been further explored recently in cardiac muscle, displaying that microtubules are integral to the hypertrophic response through the distribution of mRNA and ribosomes to the cell periphery (46). When these microtubules were disrupted, protein synthesis was observed to occur primarily in central regions and, even though occurring at ‘normal’ rates, was not able to initiate cardiac growth (46). This suggests that not only is cell periphery the primary location of protein synthesis, including for myofibrillar proteins (47), this process has to occur in this cellular location for optimal cell growth. Accordingly, our visualization of RPS6^Ser240/244^ phosphorylation in peripheral regions of human skeletal muscle extends these observations and further highlights the importance of spatial mTORC1 regulation.

In human skeletal muscle, one previous investigation has studied the cellular localization of a phosphorylated form of S6K1 (44), the upstream kinase of RPS6. The site/s probed in this particular study, threonine 421/serine 424, are not mTORC1-regulated sites as they are regulated by cyclin-dependent kinase 5 (Cdk5) (48) or c-Jun NH2-terminal kinase (JNK) (49), which means they are less associated with S6K1 kinase activity compared the mTORC1-specific site S6K1^Thr389^ (50). Nevertheless, this investigation provides information of the spatial regulation of this upstream kinase of RPS6. Here, in basal, fasted skeletal muscle, p-S6K1^Thr421/Ser424^ was found to be localized in the nuclei of skeletal muscle fibers (44), an observation we did not note in the current study (data not shown). Intriguingly, in response to an acute bout of resistance exercise, the authors observed an elevation in S6K1^Thr421/Ser424^ phosphorylation predominantly in type II fibers suggesting glycolytic fibers may respond to anabolic stimuli to a greater extent (44). However, a recent investigation demonstrated that S6K1^Thr389^ phosphorylation occurs to a greater extent in isolated type I fibers following resistance exercise and essential amino acid ingestion, particularly when presented in relation to total S6K1 protein content (51). Moreover, the phosphorylation site investigated in the current study, p-RPS6^Ser240/244^, increased to the greatest extent in type I fibers, in rodent skeletal muscle, following synergistic ablation (42). The findings presented herein somewhat agree with those reported by Goodman et al. (42) where p-RPS6^Ser240/244^ was greater in type I fibers irrespective of time point or condition. This group also recently reported elevated RPS6^Ser240/244^ phosphorylation to be greater in non-type IIB skeletal muscle fibers, compared to type IIB fibers, in mice undergoing denervation-induced atrophy. In opposition these studies however, when RPS6^Ser240/244^ phosphorylation was expressed in relation to PRE values, we did not observe an effect of fiber type on this outcome measure. This would suggest that RPS6^Ser240/244^ phosphorylation, and in principal mTORC1 activity, is perpetually greater in type I, oxidative fibers yet responds to anabolic stimuli to a similar extent in all fibers in human skeletal muscle. This elevated mTORC1 activity in oxidative fibers may contribute to the slightly greater protein synthesis rates observed in type I fibers during exercise recovery (52) and type I fiber-dominant muscles at rest and in response to exogenous amino acids (53).

Focal adhesion complexes are large, multi-protein structures which span the plasma membrane of cells in order to connect the cytoskeleton to the extracellular matrix (54) and, due to their association with costameres, transmit extracellular force to the intracellular contractile apparatus (55). Focal adhesion complexes also contain integrins and focal adhesion kinase (FAK) which can translate extracellular forces into intracellular signaling cascades to initiate cellular adaptation (56). In addition to their role in mechanotransduction, a recent *in vitro* investigation has implicated focal adhesion complexes in the activation of mTORC1 in response to amino acids and growth factors. For example, after exposure to these anabolic agents mTORC1 activity in Hela cells was seen to localize with paxillin (22), a marker of focal adhesion complexes (23, 57). Moreover, disruption of focal adhesion complexes by knockout of integral proteins or via pharmacological agents, mTORC1 could no longer be activated by growth factors or amino acids (22). Our results are consistent with an integral role for focal adhesion complexes in mTORC1 activity in human muscle as colocalization of RPS6^Ser240/244^ and Paxillin and the intensity of RPS6^Ser240/244^ staining within paxillin-positive regions were elevated at 120 min in both FED and EXFED before returning to baseline at 300 min. These two measures were conducted as colocalization calculations may be affected by changes in staining/pixel intensity across the time course, which occurs when assessing phosphorylation. Therefore, the measurement of RPS6^Ser240/244^ staining intensity within paxillin-positive regions adds further evidence of localized mTORC1 activity, confirming the colocalization data. Although trending to be greater in EXFED, the increased mTORC1 activity at focal adhesion complexes in FED as well suggests the presence of more than mechanotransduction-related signaling events in these areas. This would align with observations by Rabanal-Ruiz et al. (22) that several growth factor receptors and amino acid transporters were found to be localized within focal adhesion complexes, which have both been previously shown to contribute to mTORC1 activation. In human skeletal muscle, paxillin has also been seen to colocalize with the microvasculature (23), a location we have observed mTORC1 to translocate to following anabolic stimuli (17). Collectively, these previous data combined with our observations of localized p-RPS6^Ser240/244^ at focal adhesion complexes highlight the importance of such complexes to mTORC1 activity and cell anabolism. Interestingly, investigations in several other cell types have found dysregulated focal adhesion protein expression or localization with ageing (58, 59). Future research in older human skeletal muscle should focus on the regulation of focal adhesion complexes to understand if this contributes to the reduced ability to activate mTORC1 in this population (60, 61).

In summary, we report that RPS6^Ser240/244^ phosphorylation, a marker of mTORC1 activity, occurs predominantly in peripheral regions of human skeletal muscle fibers following anabolic stimuli. This extends our knowledge regarding the spatial regulation of mTORC1 and its contribution to mTORC1 activation in this tissue. In addition, we also show RPS6^Ser240/244^ phosphorylation is greater in type I fibers at all time points but responds to anabolic stimuli similarly in all fiber types. Finally, p-RPS6^Ser240/244^ is observed in close proximity to focal adhesion complexes, structures which contribute to mechanotransduction, growth-factor signaling and amino acid transport. These findings confirm *in vitro* data identifying focal adhesion complexes as integral contributors to mTORC1 activation and shows the preservation of this mechanism in human skeletal muscle. Future research should further focus on focal adhesion regulation to understand how these complexes may contribute to skeletal muscle dysfunction in populations who experience an inability respond to anabolic stimuli.

## Abbreviations

4EBP1: Eukaryotic translation initiation factor 4E-binding protein 1
Akt: Protein kinase B
ANOVA: analysis of variance
BSA: bovine serum albumin
Cdk5: Cyclin-dependent kinase 5
DAPI: 4′,6-diamidino-2-phenylindole
EXFED: protein-carbohydrate feeding following whole-body resistance exercise
FAK: focal adhesion kinase
FED: protein-carbohydrate feeding alone
HEK293: Human embryonic kidney 293 cells
JNK: c-Jun N-terminal kinase
MHC1: myosin heavy chain 1
MPB: muscle protein breakdown
MPS: muscle protein synthesis
mTORC1: mechanistic target of rapamycin complex 1
NGS: normal goat serum
NPB: net protein balance
p90RSK: p90 ribosomal protein S6 kinase
PBST: phosphate-buffered saline supplemented with Tween20
PRE: baseline
Rheb: ras homolog enriched in brain
RIPA: radioimmunoprecipitation assay
RPS6: ribosomal protein S6
S6K1: p70 ribosomal protein S6 kinase
SUnSET: surface sensing of translation
TBST: tris-buffered saline supplemented with Tween20
TSC1/2: Tuberous sclerosis proteins 1 and 2
v-ATPase: Vacuolar-type ATPase
WGA: wheat germ agglutinin

## Acknowledgements

The authors would like to thank Carolyn Adams, Maksym Holowaty and Hugo Fung for their assistance during data collection. The Myosin Heavy Chain 1 antibody (A4.951) developed by Dr H. Blau were obtained from the Developmental Studies Hybridoma Bank, created by the NICHD of the NIH and maintained at The University of Iowa, Department of Biology, Iowa City, IA 52242. N. Hodson is supported by a Mitacs Accelerate Postdoctoral Fellowship (IT15730). M. Mazzulla is supported by the Ontario Graduate Scholarship (OGS) program. Research was supported by a Natural Sciences and Engineering Research Council Discovery Grant (RGPIN-2015-04251) to D.R. Moore.

## Author Contributions

N. Hodson, M. Mazzulla and D.R. Moore conceived and designed research. N. Hodson and M. Mazzulla collected tissue. N. Hodson completed experimental and statistical analysis. N. Hodson and D.R. Moore interpreted results. D. Kumbhare provided medical oversight. N. Hodson drafted the manuscript. All authors edited and revised manuscript. All authors approved final version of manuscript.

